# Transformation of population code from dLGN to V1 facilitates linear decoding

**DOI:** 10.1101/826750

**Authors:** N. Alex Cayco Gajic, Séverine Durand, Michael Buice, Ramakrishnan Iyer, Clay Reid, Joel Zylberberg, Eric Shea-Brown

## Abstract

How neural populations represent sensory information, and how that representation is transformed from one brain area to another, are fundamental questions of neuroscience. The dorsolateral geniculate nucleus (dLGN) and primary visual cortex (V1) represent two distinct stages of early visual processing. Classic sparse coding theories propose that V1 neurons represent local features of images. More recent theories have argued that the visual pathway transforms visual representations to become increasingly linearly separable. To test these ideas, we simultaneously recorded the spiking activity of mouse dLGN and V1 in vivo. We find strong evidence for both sparse coding and linear separability theories. Surprisingly, the correlations between neurons in V1 (but not dLGN) were shaped as to be irrelevant for stimulus decoding, a feature which we show enables linear separability. Therefore, our results suggest that the dLGN-V1 transformation reshapes correlated variability in a manner that facilitates linear decoding while producing a sparse code.

## Introduction

A fundamental goal in neuroscience is to understand how populations of neurons represent and transform sensory information. In the mammalian visual pathway, information in the retina is relayed through the dorsal lateral geniculate nucleus (dLGN), and subsequently sent to the primary visual cortex (V1). Several hypotheses regarding how the dLGN-V1 pathway transforms visual information have been proposed. Classic sparse coding theories posit that V1 neurons encode visual information from their thalamic inputs in a sparse representation of local features (Foldiak 1990; Olshausen and Field, 1996; Olshausen and Field, 1997; Bell and Sejnowski 1997; Vinje and Gallant, 2000; Rehn and Sommer, 2006; Zylberberg et al., 2011). On the other hand, recent work inspired by machine vision has hypothesized that the visual cortical pathway transforms visual representations to make them more linearly separable in neural activity space (DiCarlo et al., 2012). Intrinsically linked to these theories is the question of how the visual population code is shaped by correlations in neural activity (Averbeck et al., 2006).

These theoretical advances have shaped our understanding of early visual processing. Multi-unit extracellular recordings have since provided evidence of sparse coding in V1 (Vinje and Gallant, 2000; Haider et al., 2010, but see Tollhurst et al., 2009). However, direct tests of whether sparse coding and linear separability theories have been limited by the difficulty of recording simultaneously from dLGN and V1 in vivo. Indeed, recent work has found that neural activity is modulated by factors such as attention and locomotion, both in dLGN (Erisken et al., 2014; Aydın et al., 2018) and in V1 (McAdams and Reid, 2005; Niell and Stryker, 2010; Polack et al., 2013; Vinck et al., 2015; Dadarlat and Stryker, 2017; Pachitariu et al., 2019). These findings complicate the comparisons of dLGN and V1 representations in different animals or recording sessions, as differences in neural activity could be caused by changes in the internal or behavioural state.

To address this, we recorded simultaneously from populations of neurons in dLGN and V1 of mice that were presented with gratings of varying spatial and temporal frequencies. Leveraging this dataset, we compared the population code in each of these areas, and how trial-to-trial spiking covariability changed the representation of visual information from dLGN to V1 within the same animal.

## Results and Discussion

### Simultaneous recordings in dLGN and V1

We used multisite silicon probes to simultaneously monitor neural populations in V1 and dLGN in mice during passive viewing of drifting gratings of varying direction and spatiotemporal frequency (Figure 1A). Mice were either awake and freely behaving (n = 3) or anesthetized (n = 1, designated separately in all figures via dotted line and/or hollow marker). V1 units had lower stimulus-evoked firing rates and were more direction selective than dLGN units (Figure 1B,C). These findings are consistent with the reduced firing and increased selectivity predicted by classic sparse coding theories (Foldiak 1990; Olshausen and Field, 1996). We next directly tested whether the dLGN-V1 transformation sparsens visual representations on a trial-by-trial basis within the same animal. To do this, we summed spikes from each neuron in time bins of varying duration (from 10 ms to 1 s), and calculated the population sparseness of V1 and dLGN activity for each trial (Vinje and Gallant, 2000). Indeed, we found that for each mouse and for all temporal windows tested, single-trial population activity of V1 units was significantly sparser than that of dLGN units (Figure 1D), and that this sparsening occurred within individual trials (Figure 1 – Supplement 1). These findings provide direct evidence of sparse coding in early vision.

**Figure 1.**
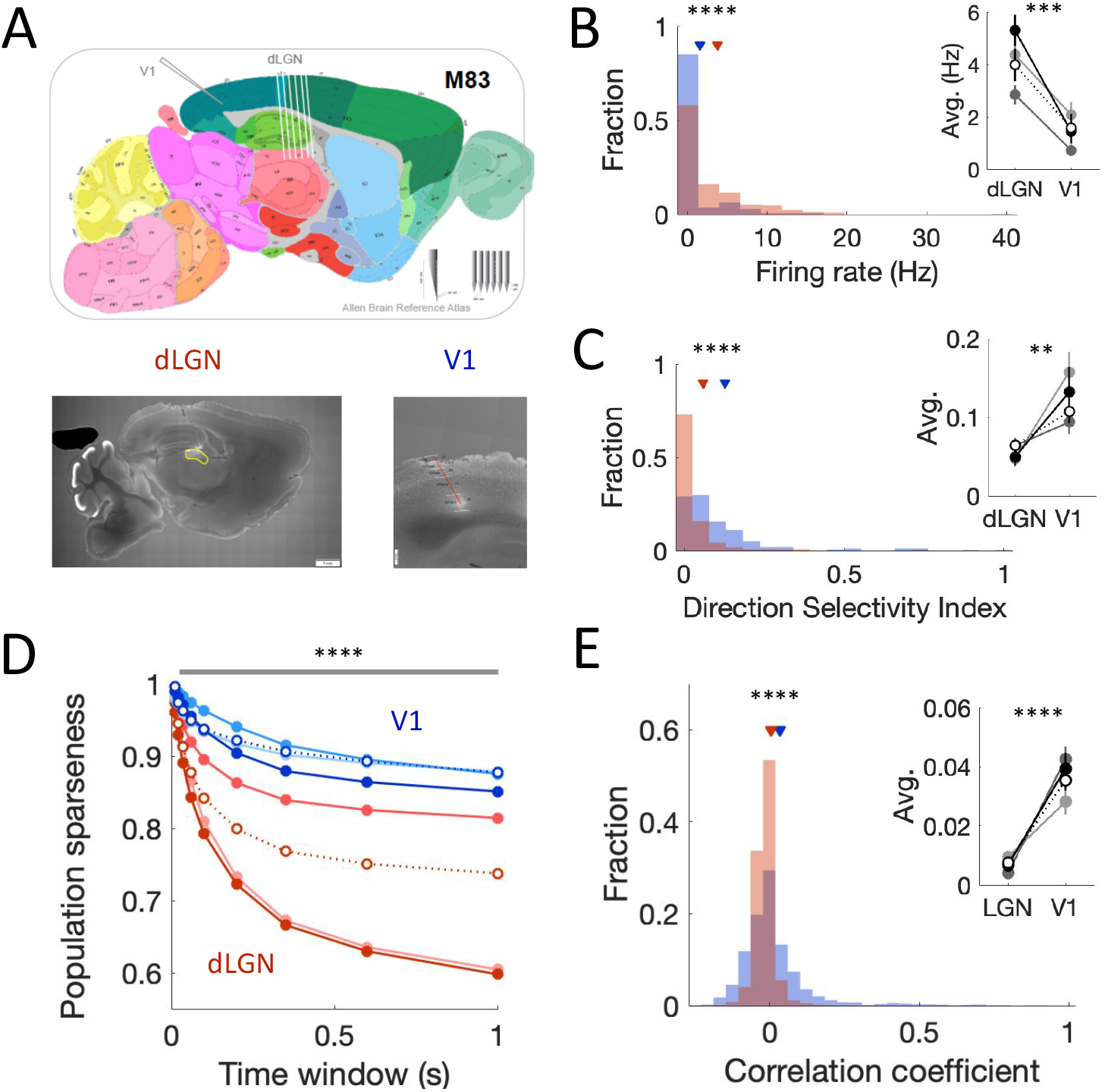
Simultaneous recordings in dLGN and V1. (**A**) Top: Typical location of electrodes in an example mouse, superimposed on Allen Brain Reference Atlas. Inset depicts single-shank and multi-shank probes used in V1 and dLGN, respectively. Bottom: Sagittal sections showing electrode tracks in dLGN (left) and V1 (right). Scale bars indicate 1 mm (left) and 200 μm (right). (**B**) Distribution of firing rates over all mice for dLGN units (blue) and V1 units (red) (p < 10^−4^, Wilcoxon rank-sum test, N = 302 dLGN units, 141 V1 units). Inset shows the change in average firing rates for each mouse separately (indicated by shade; anesthetized mouse depicted by dotted line). Stars in inset indicate minimum significance level for each mouse (p < 10^−3^ for each mouse, Wilcoxon rank-sum test). Error bars indicate SEM. (**C**) As in (B) for distribution of direction selectivity index (DSI) over all mice (p < 10^−4^, Wilcoxon rank-sum test, N = 302 dLGN units, 141 V1 units). Inset shows the change in DSI for each mouse separately (anesthetized mouse depicted by dotted line) (p < 10^−2^ for each mouse, Wilcoxon rank-sum test). (**D**) Trial-averaged population sparseness of dLGN and V1 population activity, calculated over varying time windows. Different shades indicate different mice (anesthetized mouse depicted by dotted line). Error bars indicate SEM. Stars indicate comparison of V1 vs dLGN population sparseness across individual trials (p < 10^−4^ for each mouse and for each time window tested, Wilcoxon signed rank test, N = 960 trials / stimuli). (**E**) Distribution of intraregional single-trial noise correlations over all mice (100 ms bins). V1 pairs are slightly but significantly more correlated than dLGN pairs on average (p < 10^−4^, Wilcoxon rank-sum test, N = 15,168 dLGN pairs, 2,571 V1 pairs). Inset shows the change in average single-trial noise correlation for each mouse separately (anesthetized mouse depicted by dotted line). Stars in inset indicate minimum significance level for each mouse (p < 10^−4^ for each mouse, Wilcoxon rank-sum test). Error bars indicate SEM.

To better understand how the dLGN-V1 pathway transforms activity at the population level, we next compared the correlations of spiking activity between units within the same region. Spiking correlations can loosely be categorized into signal correlations (correlations due to the stimulus tuning between neurons), and noise correlations (correlations that cannot be explained by tuning – these may reflect connectivity or modulating factors like internal state) (Averbeck et al., 2006; Cohen and Kohn, 2011; Doiron et al., 2016). We calculated spike count noise correlations in dLGN and V1 based on the joint spiking activity of pairs of neurons within single trials. Previous work has shown that nonlinear thresholding can reduce spiking correlations by quenching sub-threshold covariability of their synaptic inputs (de la Rocha et al., 2007). Therefore, naively one might expect V1 units to be significantly less correlated than dLGN units as a result of their lower firing rates. Surprisingly, we found that the average noise correlation between V1 units was slightly greater than in dLGN (Figure 1E). This difference persisted over a wide range of temporal windows (Figure 1 – Figure Supplement 2). One potential confound is that the multi-shank probe used in dLGN may record from more dispersed units. As correlations tend to decay over long distances, this could contribute to the lower dLGN correlations that we observed. However, we found the same difference in correlations when removing dLGN pairs located on different shanks (Figure 1 – figure supplement 1). In sum, we found that the the dLGN-V1 transformation sparsens population activity while slightly increasing average correlated variability.

### dLGN-V1 transformation removes impact of correlations on decoding

A significant amount of recent theoretical work has shown that correlated spiking can impact neural coding (Averbeck et al, 2006; Shamir, 2014; Kohn et al., 2016). Yet, V1 decoding studies in mice and monkeys (Panzeri et al., 2002; Averbeck and Lee, 2006; Montani et al., 2007; Poort and Roelfsema, 2009; Berens et al., 2012; de Vriees et al., 2018 -- but see Graf et al., 2011 for an exception), have generally observed very little effect of correlations on visual coding. On the other hand, previous work has shown that precisely timed spiking correlations between pairs of dLGN neurons (Alonso et al., 1996) may act as an extra information channel (Dan et al., 1998). We therefore next asked how the dLGN-V1 transformation changes stimulus decoding. In particular, we asked how the change in spiking correlations from dLGN to V1 changes how these regions represent specific visual features (e.g., direction) while remaining invariant to others (e.g., frequency).

To do this, we trained downstream decoders on either dLGN or V1 activity to discriminate between neural responses to similar visual stimuli (i.e., between responses to a pair of directions θ_1_ and θ_2_ differing by 45 degrees). We then tested the ability of these decoders to predict which of the two grating directions were presented on a single held-out trial of unknown spatiotemporal frequency (Figure 2A). For a fair comparison, V1 and dLGN populations were subsampled to the same number of neurons used for decoding in each mouse. To test the impact of correlated variability on decoding, we compared the performance of decoders based on models that captured the population activity (‘Correlated decoder’), versus those based only on the firing rates of each neuron while missing their trial-to-trial covariation (‘Independent decoder’; see **Methods**). By comparing the performance of correlated and independent decoders based on dLGN or V1 activity, we were then able to quantify how knowledge of the population correlation structure affects stimulus decoding (Figure 2B). Although overall decoder accuracies were heterogeneous across animals (Figure 2 – figure supplement 1), dLGN decoders always performed significantly better when taking into account trial-to-trial correlations (Figure 2C). In contrast, knowledge of V1 correlations had less impact on decoding (Figure 2C), despite the fact that V1 correlations were greater on average than dLGN correlations (Figure 2 – Figure supplement 2, which shows correlations for responses combined across spatiotemporal frequencies, as in our decoding task). Because the yield of our dLGN recordings was often much higher than our V1 recordings, we repeated our decoding analyses for different sizes of subsampled dLGN populations. As expected, the impact of correlations on dLGN decoding increased with population size (Figure 2D). Together, these results suggest that the dLGN-V1 transformation reshapes population activity so that correlated spiking has less impact on stimulus decoding.

**Figure 2.**
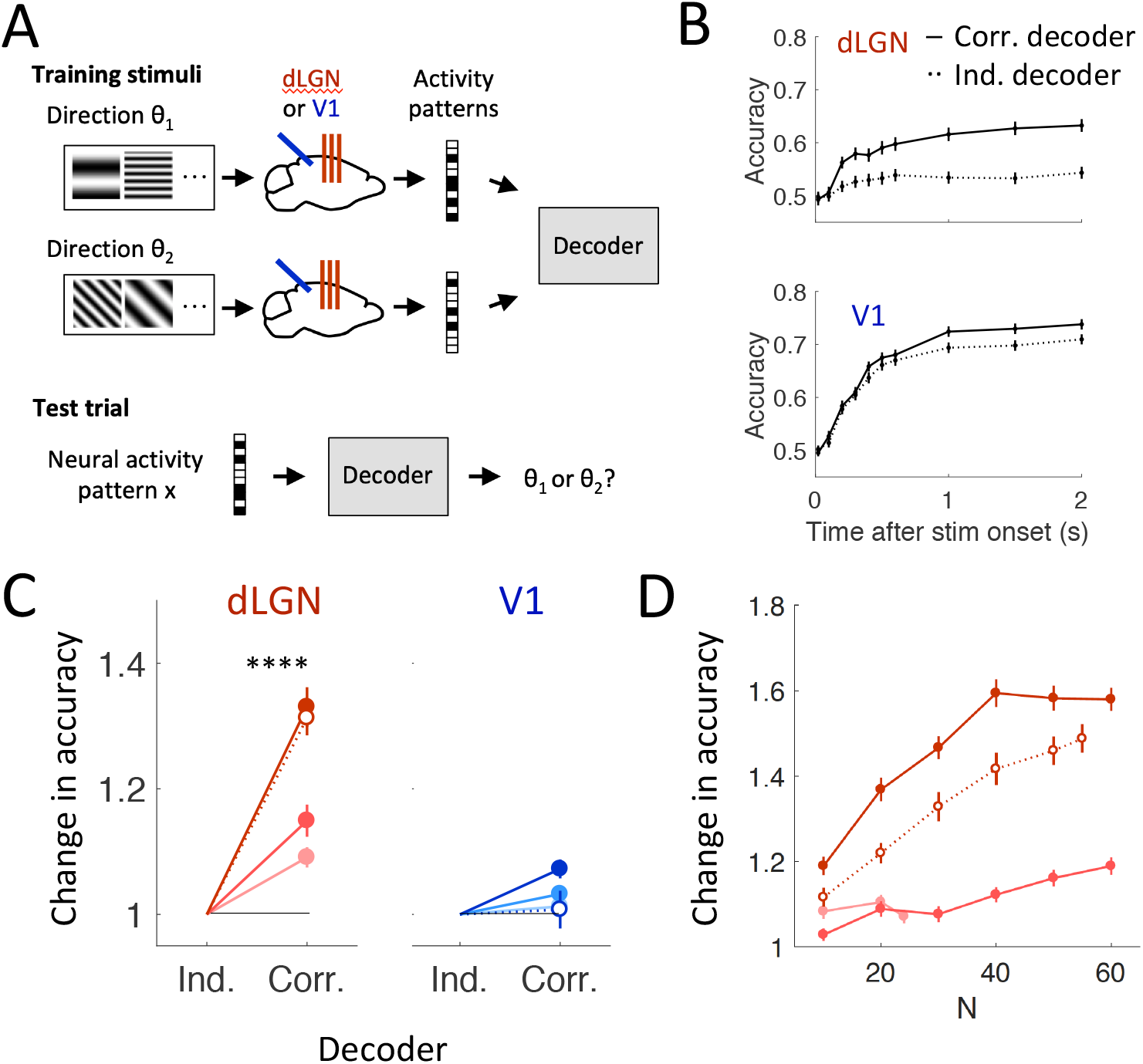
Knowledge of correlations improve decoding in dLGN but not in V1. (**A**) Schematic of decoding task. Maximum likelihood decoders were built based on the neural activity patterns of either dLGN or V1 populations in response to gratings of a given direction, while spatial and temporal frequencies were varied. These decoders were then used to predict grating direction based on the population response in a single, held-out trial (i.e., a trial that was not used for training the decoder). To compare the impact of correlations on decoding, decoders were built which either used only firing rate information (‘independent’ decoders), or used both firing rate and short-timescale (20 ms) correlated spiking (‘correlated’ decoders). (**B**) Decoding grating direction from dLGN (top) or V1 (bottom) population activity in an example mouse. Solid lines indicate the performance of a correlated-based decoder, while dotted lines indicate performance of an independent decoder. Error bars indicate SEM over 150 random samples of the held-out trial, and subsamples of the full population. (**C**) Change in decoding accuracy due to knowledge of correlated population structure, shown both for dLGN and V1 responses, and for all mice (anesthetized mouse depicted by dotted line). Knowledge of the correlated structure greatly increases decoding performance in dLGN (red, p < 10^−4^ for each mouse, Wilcoxon signed rank test, N = 150 random samples), but has a smaller impact in dLGN (blue). Error bars indicate SEM over 150 random samples of the held-out trial, and subsamples of the full population. (**D**) Change in dLGN decoding accuracy as a function of population size. Each line indicates a different mouse (anesthetized mouse depicted by dotted line). Error bars indicate SEM over 150 random samples of the held-out trial, and subsamples of the full population.

### The reduced impact of correlations from dLGN to V1 produces a more linearised population code

What is the benefit of restructuring spiking co-variability so that it has less of an impact on population decoding? Recent theories have suggested that the transformation of visual representations across successive stages of higher cortical processing helps to separate invariant representations of objects in the ventral stream (DiCarlo et al., 2012). We therefore speculated that a similar reshaping of the population code may also occur in the dLGN-V1 transformation for simpler visual features. We first showed analytically that the independent decoder can be mathematically reduced to a linear decoder (**Appendix 1**). Linear decoders are favoured because they are easy to implement in a downstream neuron, as a weighted summation and thresholding of population activity. Conversely, the correlated decoder is equivalent to a nonlinear (quadratic) decoder (Fig 3A). This fact, combined with the different effects of correlations on dLGN and V1 decoding (Figure 2C), suggests that V1 population activity may be more linearly decodable than dLGN activity. Indeed, we found that nonlinear (quadratic) decoders significantly outperformed linear decoders in dLGN, but not in V1 (Figure 3B), confirming that the dLGN-V1 transformation increases the linear separability of simple visual features.

**Figure 3.**
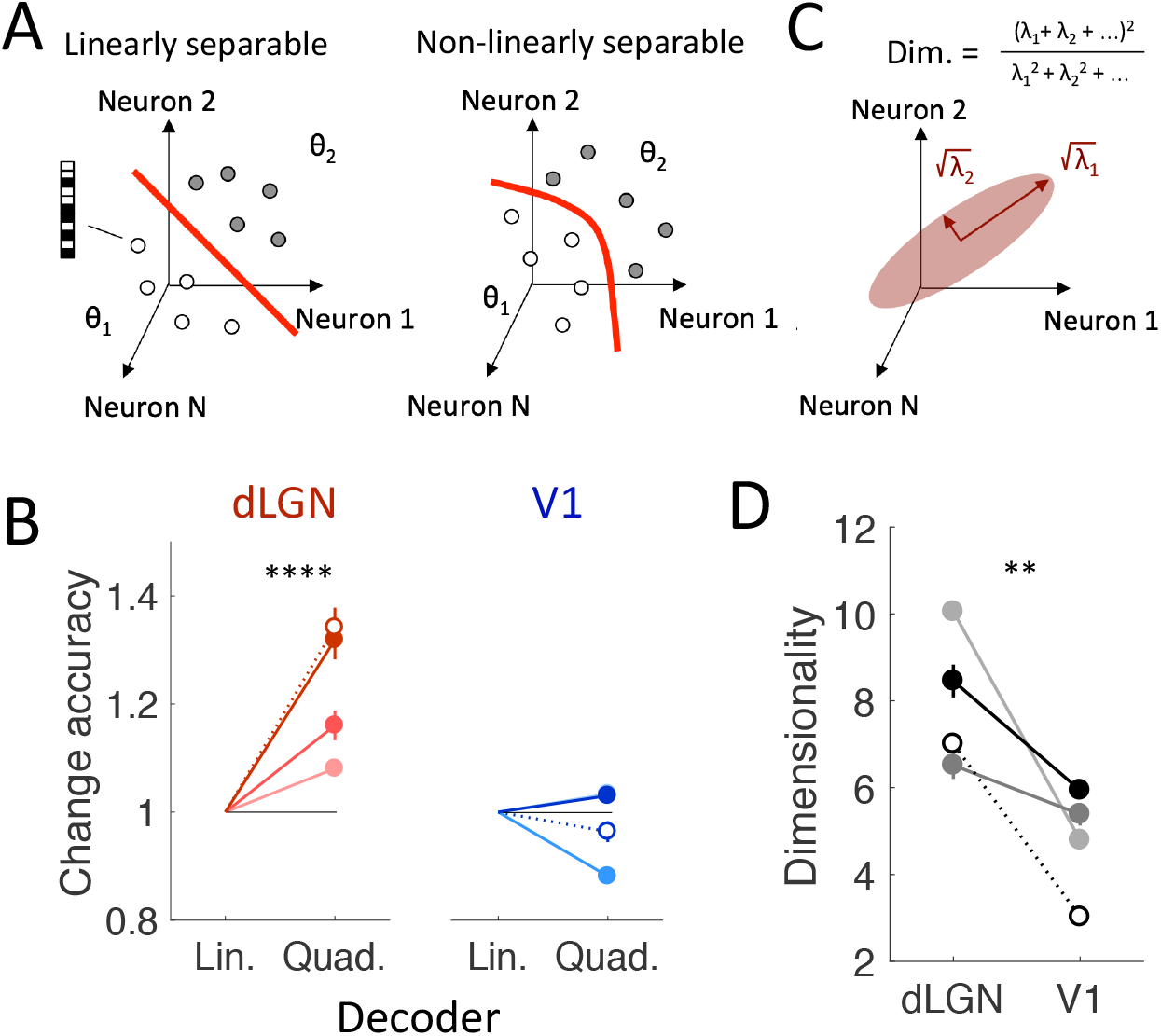
Transformation of population code makes visual information more linearly decodable. (**A**) Separability of neural representations in activity space, in which each axis represents the activity (e.g., firing rate) of a neuron. Each point represents a neural activity pattern (here, white markers represent grating stimuli of direction θ_1_, grey markers represent direction θ_2_). Left panel: In this example, the representations of the two directions can be separated by a linear separator (red). Right: In general, a non-linear separator (such as a quadratic function) is necessary to separate representations. (**B**) Change in accuracy of a quadratic decoder (compared to a linear decoder) for dLGN (red) and V1 (blue), shown for all mice (anesthetized mouse depicted by dotted line). The quadratic decoder performs significantly better than the linear decoder in dLGN (p < 10^−4^ for each mouse, Wilcoxon signed rank test, N = 150 samples), but has a smaller impact in dLGN (blue). Error bars indicate SEM over 150 random samples of the held-out trial, and subsamples of the full population. (**C**) The eigenvalues of the covariance matrix of the spiking activity are related to the extent of the neural representation along its principal directions. The intrinsic dimensionality of the neural representation can be calculated as the ‘participation ratio’ of these eigenvalues (Abbott et al., 2011; Gao and Ganguli, 2017). (**D**) Dimensionality of dLGN and V1 population activity (100 ms bins, anesthetized mouse depicted by dotted line). Population activity was randomly subsampled to 20 neurons to compare across regions / animals (p < .01 for each mouse, Wilcoxon rank-sum test, N = 30 random samples). Error bars indicate SEM.

Anatomically, the projection from dLGN to V1 is highly divergent, with hundreds of cortical neurons for every geniculate neuron in primates (Stevens, 2001). Such divergent projections are generally thought to separate neural representations by projecting them into a high-dimensional space (Cover, 1966; Albus, 1971; Rigotti et al., 2013; Babadi and Sompolinsky, 2014, Litwin-Kumar et al., 2017; Cayco Gajic et al., 2017; Cayco Gajic et al., 2019). Indeed, recent large scale imaging has found high-dimensional representations of naturalistic images in V1 population activity, consistent with this hypothesis (Stringer et al., 2019). However, we observed a decrease of the embedding dimensionality of V1 representations compared to dLGN representations (Figure 3B). One possibility is that the dLGN-V1 pathway uses different strategies – increasing dimensionality vs. reshaping representations – for complex vs. simple visual stimuli. Overall, our results suggest that the dLGN-V1 transformation can make representations of simple visual features more linearly separable by reshaping correlated spiking variability, even without increasing their dimensionality. This finding complements recent work that has shown that changes in the geometry of visual representations of objects contribute to their increased separability in deep neural networks (Cohen et al., 2019; Recanatesi et al, 2019). However, an important caveat of our interpretation is that, because we have subsampled dLGN and V1 recordings the same population size, we may miss an increase in dimensionality that could occur due to the anatomical divergence. In fact, it seems likely that these two mechanisms – increasing dimensionality and reshaping covariability – work in tandem to reshape representations of visual stimuli to be more separable. Recent advances in high density electrophysiology (Jun et al., 2017) will enable future testing of the mechanisms behind linear separability of invariant representations in large scale recordings of neural populations across the brain.

## Methods

Mice were maintained in the Allen Institute for Brain Science’s animal facility and used in accordance with the protocol approved by Allen Institute for Brain Science’s Institutional Animal Care and Use Committees.

### Extracellular multisite electrophysiology

The experimental procedure was the same as in our previously-published work (Durand et al., 2016), and summarized briefly here. Electrophysiological recordings were performed in the left hemisphere of adult C57Bl/6 mice (2–6 months, males). We performed experiments in awake (n= 3) and anesthetized mice (n= 1; urethane (1.2–1.5 g/kg, i.p.)). Using aseptic conditions, mice were first implanted with a metallic head-plate (with well) under anesthesia with isoflurane (3%–5% induction and 1.5% maintenance, 100%O2) and ketamine/xylazine (70 mg/kg, i.p.). Body temperature was maintained at 37.5°C. The head-plate was positioned and secured by Vetbond (Webster Veterinary) and Metabond (Parkell). The head-plate provides a secure way to stabilize the head and allows easy access to our targets areas (dLGN and V1). Finally, we sealed the skull surface with a thin layer of transparent Metabond and filled the well of the head-plate with Kwik Cast (WPI) until the day before the experiment, 3–8 weeks later.

The day before recordings, we performed two craniotomies under anesthesia: one directly over the dLGN and one over V1. We used two small craniotomies instead of a large one to reduce brain movement leading to probe drift. The multi-shank dLGN probes were too thin to go through the dura, so we performed a durotomy using an insulin syringe with bent tip. For reference and grounding, we inserted two screws, one over frontal cortex and the other caudal, over the cerebellum. Then, we filled the well with Kwik Cast. To allow recovery from anesthesia and surgery, the recordings were performed the next day. Awake mice went directly into the recording setup. They were head-fixed while they were free to run or remain still on a freely rotating disk. For the anesthetized mouse, we started by giving dexamethasone to avoid brain inflammation (2 mg/kg, s.c.) and atropine to keep the respiratory tract clear (0.3 mg/kg, i.p.). Then, it was head-fixed and placed on a heating pad with feedback control (ATC 1000, WPI), on a static disk. Body temperature was maintained at 37.5°C. For both awake and anesthetized recordings, the Kwik Cast was removed and the exposed cortex and skull were covered with 1% agarose in saline to prevent drying and to help maintain mechanical stability. dLGN electrodes were either a Neuronexus Buzsáki32 (32 channels with 4 shanks) or a Buzsáki64 (64 channels and 6 shanks) and were advanced vertically to a depth of 2700–2900 μm. V1 electrodes were an Edge32 (A1×32-Edge-5 mm-20–177) and were advanced to a depth 800–1000 μm. The electrodes were dipped with a lipophilic dye (DiI) allowing post hoc visualization of the electrode’s path. We then inserted the probes and used them to record brain activity. We allowed 20 minutes after implantation for the electrodes to settle, before starting the recordings. In the anesthetized mouse, the eyes were covered with a thin layer of a long-lasting lubricating and moisturizing agent (I-drop) to prevent drying. Set up and recording lasted around 5 to 6 hours.

After the recordings, mice were perfused with 4% PFA (after induction with 5% isoflurane and 1 L/min of O2). Their brains were preserved in 4% PFA, rinsed with 1X PBS the next morning, and stored at 4°C in PBS. We cut coronal or sagittal 100m sections with a vibratome. The sections were mounted using VectaShield with DAPI (Vector Laboratories) and imaged on Olympus VS-110\120 with a magnification objective of 10x to reconstruct electrode paths in V1 and dLGN (see Figure 1). Based on DiI staining, we confirmed that the probes reached dLGN (Allen Mouse Brain Atlas, Lein et al., 2007).

#### Data acquisition

Neurophysiological signals were amplified (400x), bandpass filtered (0.3-10 kHz), and acquired continuously at 20 kHz at 16-bit resolution using an Amplipex system (http://www.amplipex.com/). The spike sorting procedure was described in detail previously (Mizuseki et al., 2009). In brief, extracellular action potentials were extracted from the recorded broadband signal after high pass filtering (>800Hz) by a threshold crossing-based algorithm. Based on principal component analysis, individual spikes were automatically clustered into groups using the KlustaKwik program (Harris et al., 2000), followed by manual adjustment of the clusters using the Klusters software package (http://klusters.sourceforge.net) (Hazan et al., 2006). Only units with clear refractory periods and well-defined cluster boundaries were included in the analyses (Harris et al., 2000).

### Visual stimuli

Stimuli were generated in Python, using the Psychopy toolbox (Peirce, 2007), and were shown on an Asus PB238 screen (1920 × 1080 pixels, 51 cm wide, 60 Hz refresh rate) adjusted with a flexible arm to be 45° from the anteroposterior axis, 11 cm from the mouse’s eye, thus subtending 133° of visual field. The monitor was gamma corrected to linearize luminance. All stimuli were presented full screen.

### Gratings

We presented an extensive matrix of stimuli consisting of drifting gratings. We aimed to explore a broad range of spatial and temporal frequencies (Niell and Stryker, 2008; Andermann et al., 2011; Piscopo et al., 2013). This stimulus set consisted of gratings with eight directions (4 orientations) spaced uniformly between 0 deg and 360 deg, six spatial frequencies (0.02, 0.04, 0.08, 0.16, 0.32 and 0.64 cycles per degree), and five temporal frequencies (1, 2, 4, 8, 15 Hz) with contrast fixed at 80%. We analyzed data only from the 3 lower spatial frequencies, because the neural responses recorded at the 3 higher spatial frequencies showed unstable inter-neuronal correlations (i.e., the correlations differed substantially between different portions of the dataset, precluding accurate statistical models to be fit to the population activity).

The gratings were presented for 3 s with 1 s of mean luminance gray between successive gratings. Blank (mean gray) stimuli were randomly interleaved. Grating conditions were presented in a randomized fashion, and each condition was presented at least 7 times.

### DSI and sparseness

The direction selectivity index (DSI) of each neuron was measured following (Zylberberg et al., 2016):

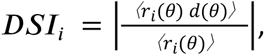

where *r*_*i*_(*θ*) is the firing rate of neuron i during all presentations of a grating with direction *θ*(regardless of spatiotemporal frequency), and *d*(*θ*) = [*cos*(*θ*), *sin*(*θ*)] is the unit vector in the direction of angle *θ*. Averages are taken over stimulus direction. Intuitively, this quantity computes the vector mean of the circularized tuning curve, normalized by the average firing rate, and takes the magnitude of the resulting vector. As a result the DSI is 1 for a neuron that only responds to stimuli of a single grating direction, and is 0 for a neuron that fires on average the same amount for any grating direction.

Population sparseness at a single time bin was quantified by (Vinje and Gallant, 2000):

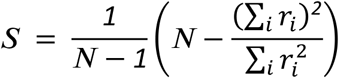

where r_i_ is the firing rate of neuron i in the corresponding time bin. This quantity measures how evenly spread spiking activity is over the population. Single-trial population sparseness was calculated by averaging S over all time bins within the 3 s trial (shown in Figure 1 – Supplement 1). The single-trial population sparseness was then averaged over all trials to obtain the trial-averaged population sparseness for each mouse (shown in Figure 1D).

### Cell selection for decoding analyses

For all decoding analyses, neurons with firing rates < 0.1 Hz were excluded due to difficulty fitting infrequent spiking statistics. Moreover, in all decoding analyses, neuronal populations were randomly subsampled so that the number of neurons used for decoding dLGN and V1 activity was the same within each mouse (Mouse M40, N = 24; M72, N = 21; M78, N = 16; M83, N = 30).

### Correlated v. independent decoding

Population decoding was assessed as two-alternative forced choice accuracy of a maximum likelihood estimation. That is, given neural activity pattern x at a single snapshot in time, the decoder chooses direction θ_1_ if P(x|θ_1_) > P(x|θ_2_). To test the impact of correlations on decoding, we compared two models of the conditional distribution P(x|θ): one that only reproduces the firing rates of the neurons (the ‘independent’ decoder), and one that also reproduces the pairwise noise correlations (the ‘correlated’ decoder). Specifically, we used maximum entropy models to fit the joint spiking activity of populations of either dLGN or V1 neurons (Gardella et al., 2019). Maximum entropy models are useful because they can fit specified statistics of the data (such as correlations and firing rates) while making no additional assumptions on the rest of the structure of the data. The maximum entropy model has the following form:

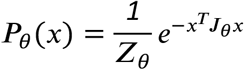

***J***_*θ*_ is a symmetric matrix of pairwise interaction terms capturing the coupling between neurons, and ***Z***_*θ*_ is a normalizing factor. Spiking activity x was binned in short 20 ms time windows. For parameter fitting, we used Minimum Probability Flow (Sohl-Dickstein et al., 2011). For correlated decoders the full matrix ***J***_*θ*_ was fit to match the firing rates and pairwise correlations, while for independent decoders ***J***_*θ*_ was constrained to be diagonal (and was fit to match only the firing rates). ***Z***_*θ*_ was approximated using the Good-Turing estimate (Haslinger et al., 2013). Note that a separate model was fit for each grating direction. One trial per orientation was held out for testing the model. This process was repeated 150 times with different choices of randomly subsampled populations and held-out trials. Upon testing, multiple timepoints were treated as statistically independent to obtain decoding accuracy as a function of time throughout the held-out trial (e.g., Figure 2B). Finally, decoding accuracy was averaged over all pairs of grating directions differing by 45 degrees. Similar results were obtained when averaging over all possible pairs of directions.

### Linear vs. quadratic decoding

Linear separability was determined using cross-validated logistic regression to discriminate between pairs of stimuli. For each pair of grating directions, a linear classifier was built by fitting a generalized linear model (GLM) with a logit link and binomial distribution to classifiy the vector of 20 ms binned, binarized neural activity. For quadratic decoders, regressors additionally included cross-terms (i.e., *x*_*i*_*x*_*j*_). Linear (or nonlinear) decoding accuracy was then calculated by predicting the stimulus based on the population activity averaged over the first 600 ms of a held-out test trial. Decoding accuracy was averaged over all pairs of grating direction and over 150 random choices of subsampled populations and held-out test trials.

### Dimensionality

To compare the dimensionality across mice and across regions, all dLGN and V1 populations were randomly subsampled to 20 neurons. The dimensionality was then measured as the eigenvalue participation ratio (Gao et al., 2017; Litwin-Kumar et al., 2017):

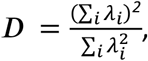

where each *λ*_*i*_ is the ith eigenvalue of the covariance matrix of the 20 ms binned spiking activity over the first 600 ms following stimulus presentation. This can be viewed as a generalization of dimensionality (as typically measured as the number of principal components required to capture a fixed percentage of the variance) to continuous values.

## Code availability

All analysis code will be made available upon publication at: https://github.com/caycogajic/LGN-V1-neural-code

## Acknowledgements

We thank the many staff members of the Allen Institute and especially the In Vivo Sciences team for surgeries, the Engineering team, the Imaging Team and Naveen Ouellette for expert assistance; and Allen Institute founder, Paul G. Allen, for his vision, encouragement and support. JZ is an Associate Fellow of CIFAR, in the Learning in Machines and Brain Program, and gratefully acknowledges the following funding sources: Sloan Fellowship; Canada Research Chairs program; NSF-DMS Grant 1056125, and National Science and Engineering Research Council of Canada (NSERC).

## Appendix 1

Suppose we have M neural activity patterns 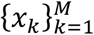, with each pattern *x*_*k*_ ∈ [0,1]^*N*^ from an unknown stimulus Θ. Under the maximum entropy model, the probability of each sample *x* is given by:

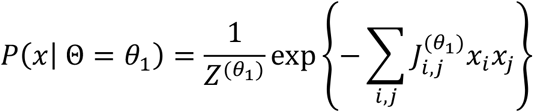

We estimate decoding accuracy using the maximum likelihood estimate based on the conditional probabilities *P*(*x*|Θ = *θ*_1_ in a two-alternative forced choice task. Multiple patterns (which are taken as sequential binned activity patterns during the held out task) are treated as independent by the decoder.

Specifically, the decoder guesses stimulus *θ*_1_ over stimulus *θ*_2_ if *P*({*x*_*k*_}|Θ = *θ*_1_ > *P*({*x*_*k*_}|Θ = *θ*_2_. Assuming the samples are independent, this reduces to the following:

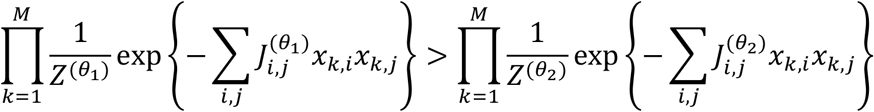

Taking the log of both sides:

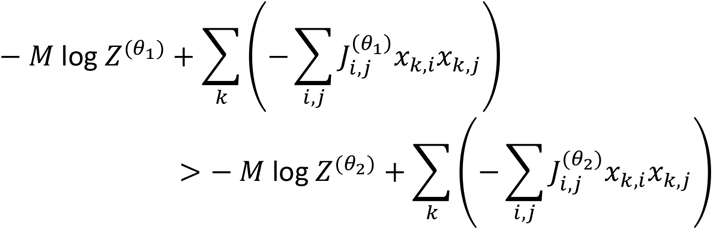

Simplifying this equation yields:

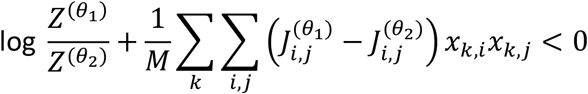

For a correlated decoder (pairwise maximum entropy model), this is equivalent to the following quadratic separator:

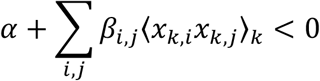

where 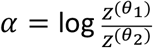 and 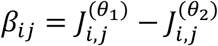. For an independent decoder, J is diagonal therefore all cross terms vanish. As *x* is binary (hence *x*^2^ = *x*), this reduces to a linear separator of the time-averaged activity of single units:

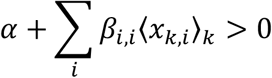

Therefore, the correlated decoder is equivalent to a quadratic decoder, and the independent decoder is equivalent to a linear decoder.

---

**Figure 1 – figure supplement 1.**
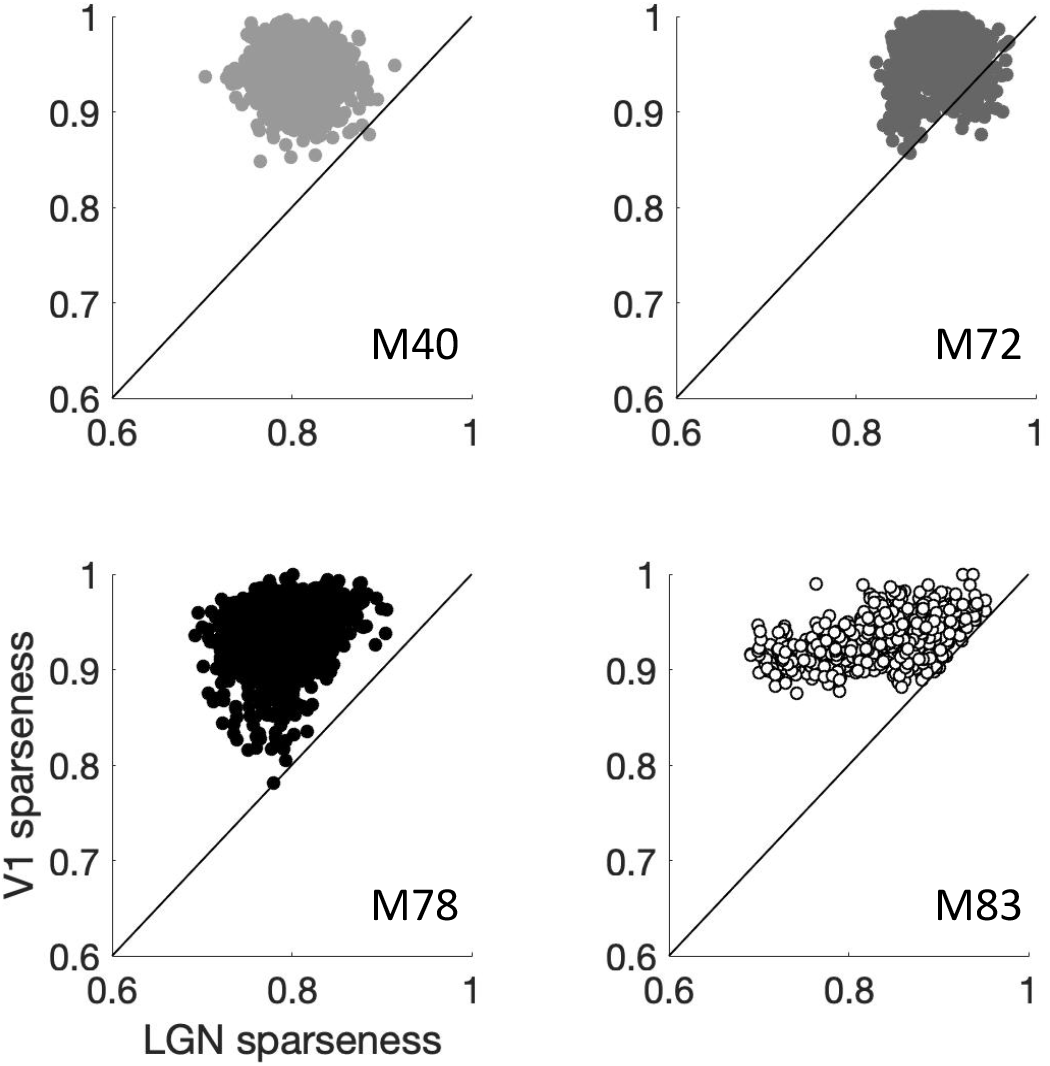
dLGN vs V1 population sparseness on single trials, calculated for 100 ms binned responses. Each point indicates a different direction and spatiotemporal frequency and trial, and each panel depicts results for a different mouse (M83 indicates anesthetized mouse).

**Figure 1 – figure supplement 2.**
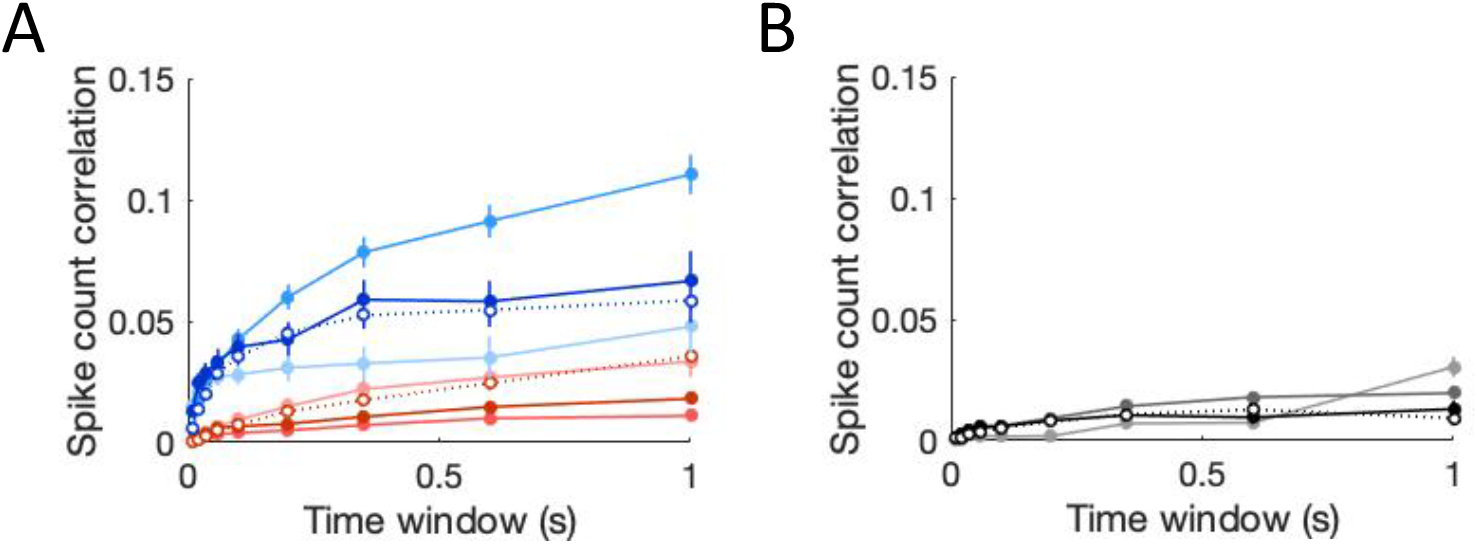
Intraregional correlations are smaller than interregional correlations. (**A**) Average intraregional correlations between dLGN pairs (red) or V1 pairs (blue), calculated over varying time windows. Error bars indicate SEM. (**B**) Same as (A), but for interregional correlations between dLGN and V1 units.

**Figure 1 – figure supplement 3.**
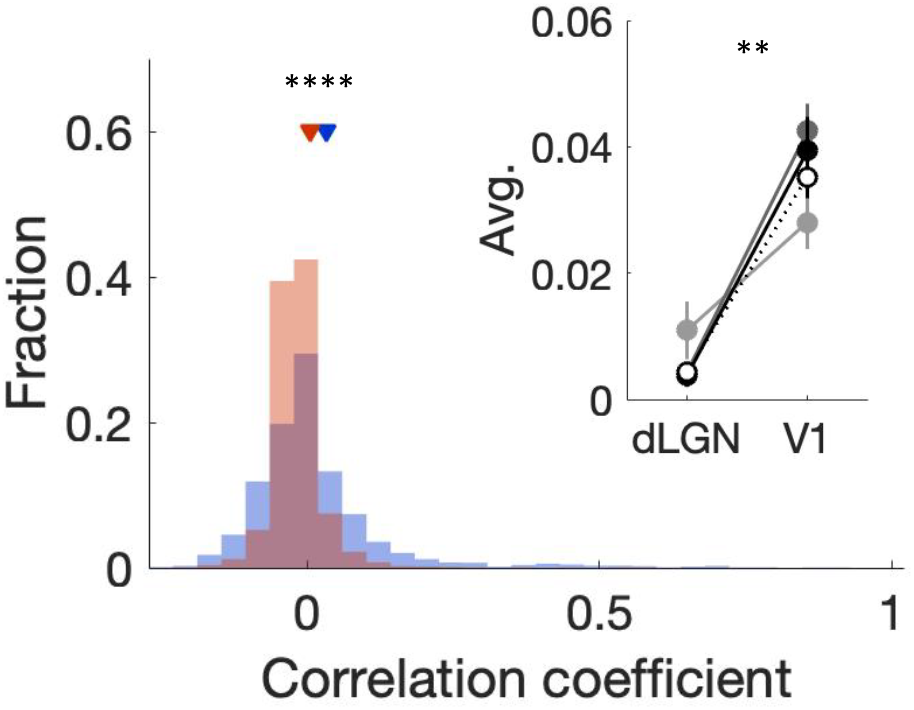
Distribution of noise correlations separated by probe shank. Red histogram now shows the distribution of correlations between dLGN units on the same shank. V1 distribution (blue) is identical to data shown in Figure 1E. V1 pairs are slightly more correlated than dLGN pairs on average (p < 10^−4^, Wilcoxon rank-sum test, N = 2767 dLGN pairs, 2571 V1 pairs). Inset shows the change in average noise correlation for each mouse separately. Stars indicate minimum significance level for each mouse (p < .01 for each mouse, Wilcoxon rank-sum test). Error bars indicate SEM.

**Figure 2 – figure supplement 1.**
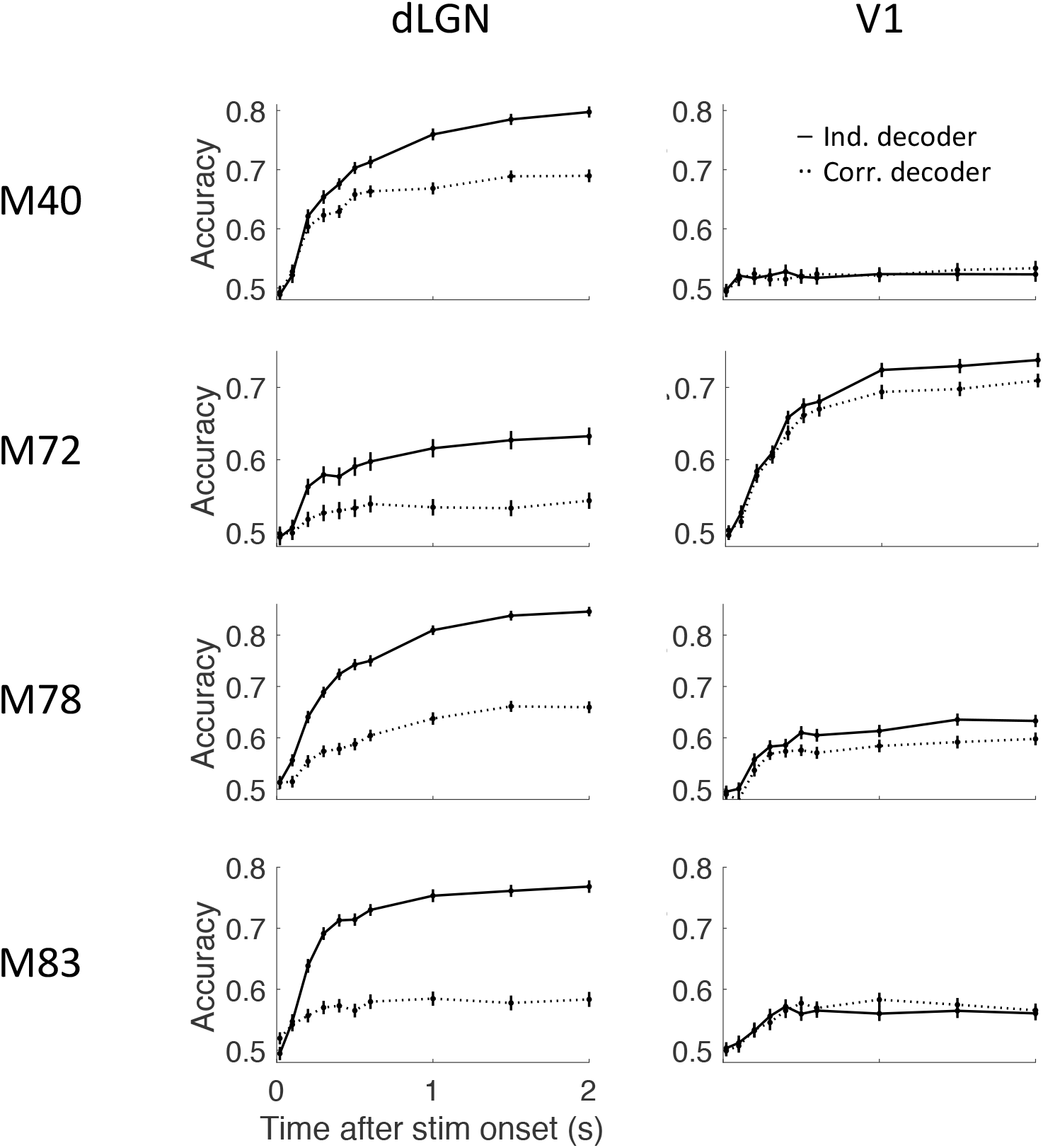
Decoding accuracy for each region and each mouse (M83 indicates anesthetized mouse). Each panel as in Figure 2B.

**Figure 2 – figure supplement 2.**
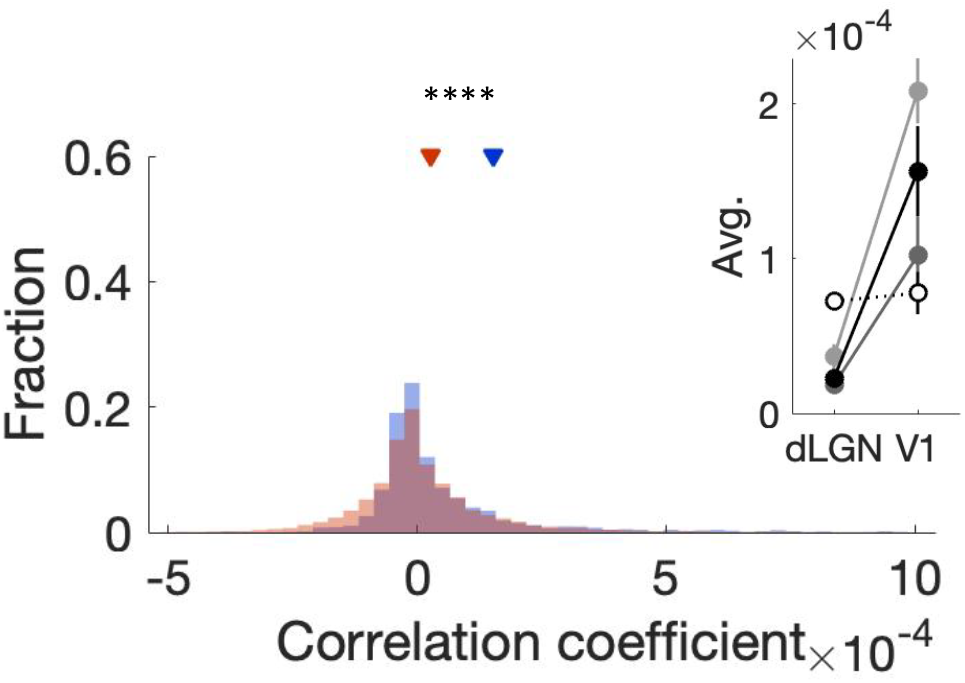
Noise correlations of decoder, that is, correlations of 20 ms binned dLGN or V1 spiking activity, calculated over all trials in which the visual stimulus has the same grating direction. As for the single-trial noise correlations (Figure 1E), dLGN correlations are lower on average than V1 correlations (p < 10^−4^, Wilcoxon rank-sum test). Inset shows change in average correlation separately for each mouse. Note that, unlike Figure 1E, the difference between dLGN and V1 correlations is not statistically significant for each mouse.

